# Model-based design and analysis of life table experiments for insect vectors

**DOI:** 10.1101/2020.03.05.978684

**Authors:** Kamil Erguler, Murat Can Demirok, Filiz Gunay, Mina Petrić, Mihaela Kavran, Dušan Veljko Petrić, Bulent Alten

## Abstract

Life tables help identify physiological differences in distinct development stages and detect potential vulnerabilities for conservation and control. However, cataloguing mortality, development, and fecundity by following each individual could be challenging in insects due to interweaving generations and development stages.

Here, we propose to use age- and stage-structured population dynamics modelling to help derive life table characteristics from the observed dynamics of reared populations. We examine a hypothetical case, a simulated population with known life parameters, and two experimental cases, observations of the population dynamics of the mosquito vectors *Culex quinquefasciatus* and *Culex pipiens*, to demonstrate that model-based inference can correctly identify life parameters from longitudinal observations. The analysis reveals not only the differential physiological behaviour of distinct development stages, but also identifies the degree to which each parameter can be inferred from the data.

The methods introduced constitute a model-based approach to identifying life table characteristics from incomplete longitudinal data, and help to improve the design of life table experiments. The approach is readily applicable to the development of climate- and environment-driven population dynamics models for important vectors of disease.

## Introduction

Life tables provide valuable insights into the dynamics and environmental dependence of populations [3, 28]. They have been used extensively for species conservation by identifying sensitive life stages [44] and key environmental drivers [51], and for designing effective control strategies against pests and disease vectors [3, 38, 40]. Life tables have become an essential component of mathematical models representing population dynamics by providing the biological foundations for model development [17, 41, 50].

A common practice of constructing life tables involves following a cohort of individuals in time and cataloguing a set of key life processes including mortality, development, and fecundity [3]. Typically, factors such as age-specific survivorship (the number of females or males on any day) are recorded and net reproductive rate (the expected number of females produced by the females of a cohort) is estimated. However, the list is adapted to fit the special needs of the species being examined [2].

Mortality is a key component of life tables; however, for many insect vectors, including mosquitoes, ticks, and small biting flies, it is difficult to assess the viability of certain stages until development commences to the next stage. For instance, egg viability is commonly quantified through larval hatching, *i.e.* eggs which do not develop into larvae after an appropriate amount of time are assumed dead [4, 19, 37].

Often, subsequent development stages overlap resulting in variability being amplified as development progresses. It might be impractical to isolate individuals or technically impossible to tag and follow each individual throughout development. Such interventions could impact development and survival, and result in inaccurate derivations [20, 47]. Multiple cohorts could be constructed to study the characteristics of different stages; however, this could restrict the experimental setup [12, 24].

Mathematical models have long been used to study and improve the understanding of many physical, chemical, and biological processes [5, 13]. By definition, a model can only represent a simplified version of the reality, and it is informed by prior knowledge, which comprises past observations, relevant literature, and expert opinion [36]. Canonical modelling approaches capture population heterogeneity partially by incorporating discernible development stages as distinct entities, *i.e.* stage-structured models [11, 27, 31]. Only a subset incorporates time-delay to represent time-driven heterogeneity as a result of, for instance, differential development rates [29, 39, 42]. Age- and stage-structured population dynamics modelling offers an alternative to represent heterogeneity in natural populations by propagating age-structure in distinct development stages [14].

The age-structured discrete-time population dynamics model (sPop) [14] is an extension of the works of Leslie [32] and Lefkovitch [11, 31]. As specified by the model, individuals are grouped into distinct stage classes but retain their developmental ages, *i.e.* the level of completion of their development. Thus, individuals in each age-development group are subject to the same survival, developmental, and reproductive characteristics. The model requires programmatic handling of a dynamic set of difference equations, and this is provided by the sPop package implemented in C, Python, and R [14]. The approach was first introduced by Erguler *et al.* (2016) [17] for modelling the environment-driven population dynamics of *Aedes albopictus*. Erguler *et al.* demonstrated in subsequent work that age- and stage-structured population models can be useful in representing field populations and disease transmission [15, 16]. This is achieved by inferring appropriate life table characteristics with regard to field observations and large-scale meteorological datasets in the absence of microenvironment conditions.

In this study, we elaborate on the model-based inference approach to identify life table characteristics under controlled laboratory settings. We follow common experimental practices where observations of viability and stage transformations are recorded at regular time intervals to the extent possible. We investigate the development of typical multivoltine insects with constantly overlapping generations and aim to derive parameters pertaining to life table characteristics at different stages. We propose that mathematical models and methods of inference can be used to derive these parameters and to relax, to a certain extent, the constraints on developing more informative but complex study designs.

Although the approach is generally applicable to any population with age-dependent heterogeneity, we focus on two important mosquito vectors, *Culex quinquefasciatus* and *Culex pipiens*, with complex dependence on climate and environmental factors. We aim to develop an initial model-based understanding of these vectors, and establish a foundation for developing environment-driven population dynamics models to help predict their seasonality and abundance.

## Methods

We propose a model-based inference protocol, which involves model construction, parameter optimisation, and posterior sampling. The process begins with the construction of a representative model structure, which relies on a set of parameters to be configured. Each configuration is a proposal and can be conceptualised as a point in a multidimensional space. The dimension of this space is given by the total number of parameters employed by the model. The process continues with the estimation of the most suitable configuration by using an optimisation algorithm, and then, the sampling of a set of suitable points from around the optimum using approximate Bayesian computation.

In the Bayesian context, the prior probability of a particular parameter configuration is informed by expert opinion and the existing knowledge base. The posterior probability, on the other hand, is determined by the ability of the model-parameter combination to replicate new observations. According to this,

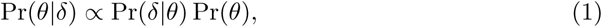

where Pr(*θ*) is the prior probability of the parameter configuration *θ* and Pr(*δ*|*θ*) is the likelihood, *i.e.* the probability of making the observation *δ* by simulating the model with *θ* [30, 48].

The prior probabilities can be the same among all proposals or set in favour of a particular hypothesis. The likelihood, however, is not often straightforward to define or evaluate. Deterministic models require explicit accounts of observational/intrinsic noise and time-correlation, which might be evident in the data. Although stochastic models can account for the noise, the estimation of the likelihood is, in most cases, computationally expensive. In response, approximate methods have been developed [35, 49]. In approximate Bayesian computation (ABC), likelihood is replaced with a simulation-based distance function, *f*, and the posterior is approximated with a given threshold *ϵ*. As a result,

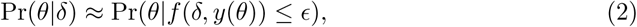

which tends to Pr(*θ*|*δ*) as *ϵ* → 0. Here, *f* is the distance between the data and the corresponding model output *y*(*θ*).

Despite the use of approximation, exploring the entire posterior distribution is often computationally demanding, particularly when the number of parameters is large [34]. In order to reduce the computational demand, we propose generating posterior samples around the optimum posterior parameter. In our previous work, we referred to this partial posterior sample as the posterior mode and demonstrated that it could be useful for model development especially when competing models are complex [15–17].

A posterior mode, Θ, is defined as a model-parameter combination which includes a subset of all possible parameter configurations [17]. In the context of ABC, the approximate marginal probability of a posterior mode can be derived from Eqns. 1 and 2 as

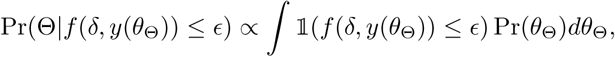

where *θ*_Θ_ represents a parameter configuration from Θ and 𝟙 represents an indicator function returning 1 if *f*(*δ, y*(*θ*_Θ_)) ≤ *ϵ* and 0 otherwise.

In order to identify the optimum value of the posterior, we propose using an optimisation algorithm on the likelihood, if available, or on the distance function. Due to its simplicity and broad applicability, including deterministic and stochastic contexts, here, we employed the Metropolis-Hastings Markov chain Monte Carlo (MCMC) algorithm [7, 26]. In S1 Text, we present a sample R script to perform parameter optimisation with MCMC.

Once an optimum parameter configuration is identified, sampling can be performed using any posterior sampling algorithm, such as the canonical rejection method or MCMC [6]. Toni *et al.* (2008) developed a sequential Monte Carlo algorithm (ABC-SMC) to sample from the approximate posterior distribution [49]. ABC-SMC is powerful, widely applicable, and straightforward to implement.

Here, we adapted the ABC-SMC algorithm to yield samples predominantly from a posterior mode. We achieved this by initiating the algorithm at an optimum parameter configuration and executing it with a fixed distance threshold. Eventually, a stationary distribution emerges which represents a sample from the posterior mode around the optimum. It is possible to obtain such a sample from a single MCMC chain originating from the same optimum provided that the chain is sufficiently iterated. However, the ABC-SMC algorithm provides a computationally feasible alternative [49]. In S2 Text we present a sample script to run the algorithm in R.

## Results

In order to demonstrate the added value of the model-based inference protocol, we examine a hypothetical scenario and two experimental case studies on important mosquito vectors.

### Case 1: Inference with a known model

In this section, we consider that the model and its parameter configuration are known. The data include noise arising from the stochasticity of the model but no observational error. We demonstrate the process described in Methods, and assess the performance of four representative modelling approaches in describing the system.

#### Target model: stochastic sPop model

We hypothesised a typical life table experiment which starts with a given number of mosquito eggs, and the eggs are observed to develop gradually into larva and pupa. In order to represent this experiment, we used a stochastic sPop model with fixed daily survival and gamma-distributed development times. Through this model, we assumed that development rate is time-dependent, *i.e.* the probability of completion increases gradually as time passes, and the individuals at the same developmental age can be grouped together.

We present in S3 Text an implementation of this model in R. The following is a set of stochastic difference equations describing the daily survival and development of eggs, larvae, and pupae.

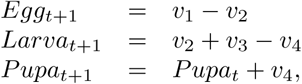

where

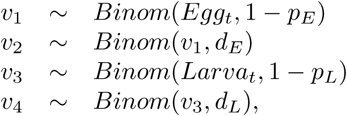

and *p* and *d* represent the daily probabilities of mortality and development, respectively. The model implies that eggs and larvae survive at fixed daily probabilities (1 − *p*), egg and larva development occurs with a daily probability of *d*, given survival, and larvae accumulate as pupae when they complete development. While survival is independent of age, the daily development probability is age-dependent and follows a gamma distribution with a given mean (*µ*_*d*_) and standard deviation (*σ*_*d*_).

In Figure 1 we present three realisations of this model simulated with an initial condition (*x*_0_, *y*_0_, *z*_0_) = (100, 0, 0) and an arbitrary set of parameter values (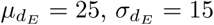, *p*_*L*_ = 0.05, 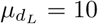, and 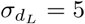). In the context of this hypothetical case, these model parameters can be regarded as life table parameters. In addition to the sPop model, we selected three modelling approaches based on their frequency of encounter in life sciences — more specifically, in ecology and epidemiology [5] — and proposed potential explanations for the data in Figure 1. We investigated each one separately and examined their advantages and disadvantages in identifying life table parameters. We employed a generic score function, the sum of squared errors (SS), as the distance function.

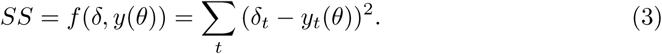

**Figure 1.**
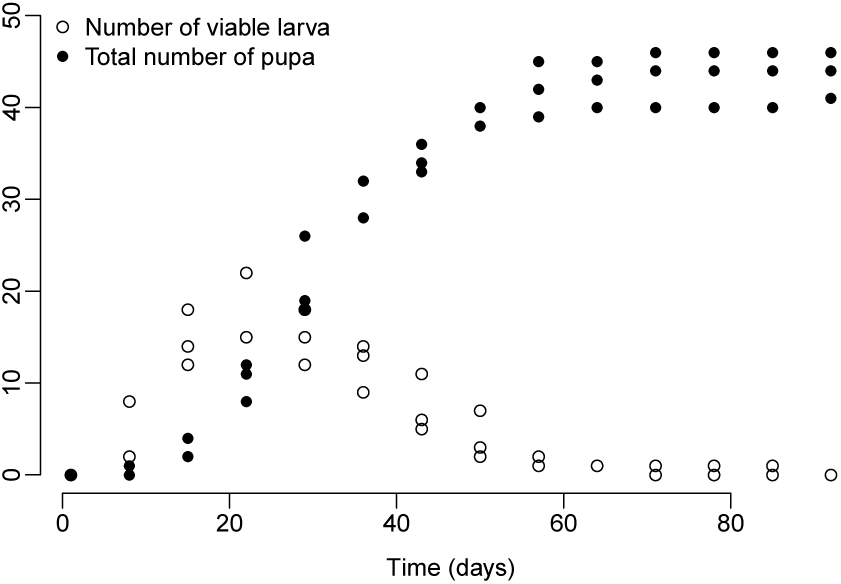
Simulated data of a typical life table experiment. The number of viable larvae (hollow circles) and the total number of pupa emergence (solid circles) are shown for each day. Three replicates were simulated starting with 100 eggs using the stochastic sPop model as described in the main text.

#### Proposed model 1: age- and stage-structured difference equations model (sPop)

To begin with, we proposed a deterministic sPop model representing the egg and larva stages with fixed daily survival and gamma-distributed development times. The products of developed larvae are accumulated in the third stage as pupa similar to the stochastic version explained above.

The difference equations describing the model are given below, while a typical implementation of the model in R is given in S4 Text.

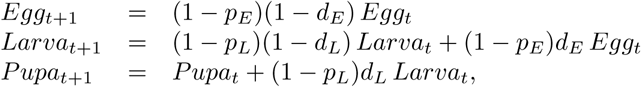

where *p* and *d* represent daily fraction of mortality and development, respectively. The model employs 6 parameters in total: *p*_*E*_, 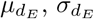, *p*_*L*_, 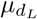, and 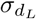, where *µ* and *σ* represent the mean and standard deviation, respectively, of the corresponding gamma-distributed development times.

### Proposed model 2: stage-structured difference equations model (Diff)

Lefkovitch in 1965 [31] introduced a generic stage-structured population dynamics model by grouping individuals into distinct stage classes with identical life table characteristics. In a modelling study for loggerhead turtles, Crouse *et al.* (1987) described a method for estimating the projection matrix, which quantifies daily change in each stage and between stages, using life tables [11].

The model, which is referred to as the Diff model in this context, permits a heterogenous stage class where the age of individuals are distributed as a function of survival probability (*q*) and development time (*d*); however, instead of iterating this distribution, which is essentially done in sPop, daily mortality and transition rates are estimated for a stage by aggregating the distribution.

In essence, the following rates are estimated:

- The daily fraction of individuals surviving and growing into the next stage:

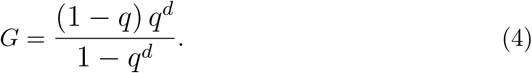
- The daily fraction of individuals surviving and remaining in the same stage:

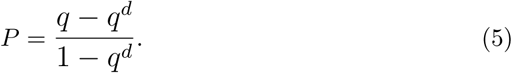

The deterministic difference equations describing the Diff model are given below, and a typical implementation of the model in R is presented in S5 Text.

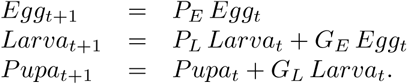

The model represents eggs progressing into larvae, which subsequently develop into, and accumulate as, pupae. In total, the Diff model employs 4 parameters: *p*_*E*_, *d*_*E*_, *p*_*L*_, and *d*_*L*_, where mortality (*p* = 1 − *q*) is replaced with survival in concert with the sPop model.

#### Proposed model 3: ordinary differential equations model (ODE)

The use of ordinary differential equations (ODEs) in vector population and disease transmission modelling dates back to the early twentieth century [43, 45]. More recently, ODE models have been developed to represent climate-driven population dynamics of many vector species [8, 50].

Despite their prevalence and relative ease of use, ODE models poorly represent population heterogeneity and the time-delay caused by underlying biological processes. Nevertheless, we followed the established practices and developed a deterministic population dynamics model to represent the transition of eggs to larvae, and then, to pupae.

The equations describing the ODE model are given below, and a typical implementation of the model in R using the deSolve package is presented in S6 Text.

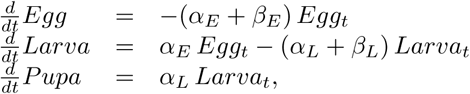

where *α* and *β* represent instantaneous development and mortality rates, respectively. In total, the model employs 4 parameters: *α*_*ϵ*_, *β*_*ϵ*_, *α*_*L*_, and *β*_*L*_.

#### Proposed model 4: delay differential equations model (DDE)

Nisbet, Gurney, and Lawton developed a stage-structured delay differential equations (DDEs) model to represent insect population dynamics [25, 39]. The authors employed instantaneous rates of mortality and fecundity, and incorporated time-delay in the development process. Recently, Kettle *et al.* implemented the model in R, which is available in the stagePop package [29].

The model enables dynamic variation in development times; however, it assumes that all individuals in a stage behave in exactly the same way regarding survival and development. This implies a homogenous stage where a sharp transition to the next stage accrues from a cohort of individuals. As a potential workaround for representing developmental variation in a stage, the authors proposed inventing a plausible population history for some time before zero until the required (or observed) variation builds up [25, 39].

Here, we employed a parameter, *τ*_*t*_, to control the rate of entry to the initial stage. In this context, variable entry times can be seen as variable starting times for development. In turn, one observes a seemingly gradual completion of development and a smooth transition between subsequent stages. Although the authors recommended generating the population history some time before *t* = 0, in this context, variable entry would yield a similar behaviour but a shift in development times. For instance, the minimum and maximum development times for variable entry are *t*_*d*_ − *τ*_*t*_ and *t*_*d*_, respectively, where *t*_*d*_ is the time it takes to the first completion of development. Conversely, the minimum and maximum development times for the population history approach are *t*_*d*_ and *t*_*d*_ + *τ*_*t*_, respectively.

The consequence of either of the approaches is that each individual follows exactly the same development process, but from a different time frame. At any instant, while some individuals are commencing development, others might have already finished it. Development always takes the same time. In cases of population interventions, such as blood feeding, or when different stages must have different developmental age distributions, this workaround may not be applicable. Nevertheless, in this context where only two stages are of interest, the approach seems applicable.

In order to represent a desired distribution of heterogeneity, *e.g.* gamma, negative binomial, or Gompertz [21], a customised rate of entry can be constructed from the derivative of the distribution function. A simple first-order approximation is to assume that individuals arrive at a constant rate, which results in a uniformly distributed transition time. Here, we adopted a constant rate of entry to examine its impact on the model and fit to the data.

A typical implementation of the DDE model in R using the stagePop package is given in S7 Text. The equations describing the model can be written as

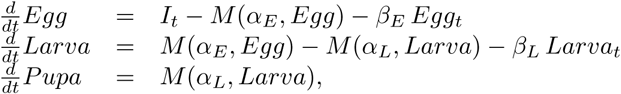

where *α* and *β* represent per capita development and death rates, respectively, and *M* is a function of *α* and the historical state of the corresponding stage. In this equation, *I*_*t*_ represents the rate of entry to the egg stage, and it is given by

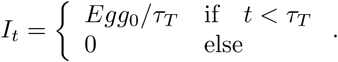

where *Egg*_0_ is the desired size of the initial egg population and *τ*_*T*_ is the time allocated to reach *Egg*_0_. Consequently, *τ*_*T*_ controls the variability of development in the egg population. In total, the model has 5 parameters: *α*_*E*_, *β*_*E*_, *α*_*L*_, *β*_*L*_, and *τ*_*T*_.

#### Assessment of model performance

As a result of the optimisation-posterior sampling procedure, described in Methods, we observed that the four models offered four different explanations for the data. We list the distance thresholds (*ϵ*) and minimum distances achieved by each model in Table 1. In Figure 2, we present the output ranges of each model in comparison with the observations.

**Table 1.** Model performances against the simulated life table experiment.

| Model | Threshold ($\epsilon$) | Min. distance |
| --- | --- | --- |
| sPop | 362.471 | 331.4 |
| Diff | 1041.56 | 1031 |
| ODE | 1118.7 | 1048 |
| DDE | 558.181 | 535.8 |

**Figure 2.**
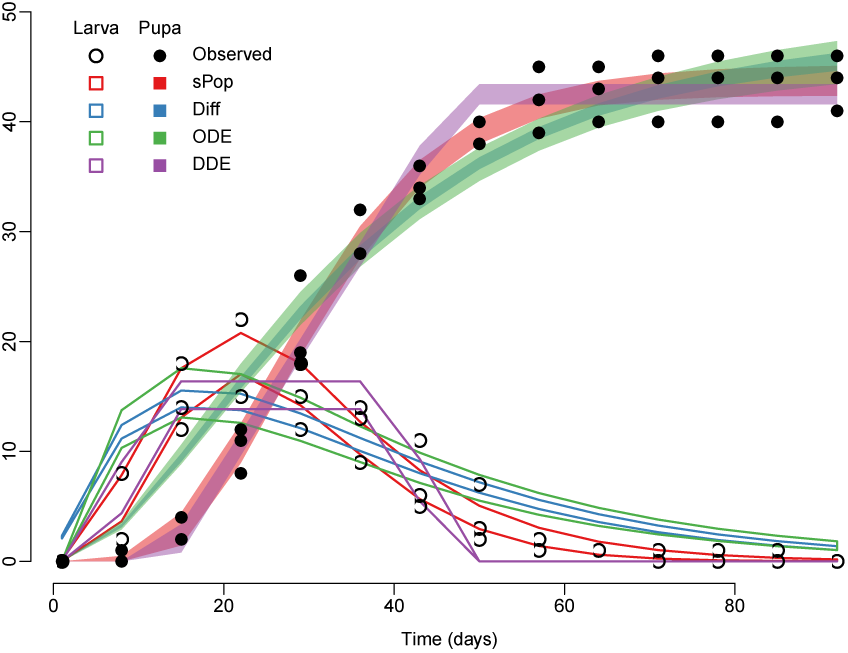
Comparison between the observations and model fits. Shaded and outlined trajectories display the ranges of pupae and larvae, respectively, as simulated using one of the four models and its corresponding 1000 posterior parameter samples. The observations are shown as points as in Figure 1.

As a result, we observed that the sPop model performed better than the other three in offering an explanation to the observations. The sPop and target models exhibit significant similarities; their main difference is the omission of stochasticity in the former. As a consequence, the observed performance of the model was expected.

As the second best explanation, the DDE model achieved distances comparable to those of sPop, and performed far better than the Diff and ODE models. The results suggest that the uniform distribution in entry times, employed for the DDE model, contributed to explaining the observed gradient in stage transition.

Finally, we observed that the Diff and ODE models failed to represent the time delay emerging from development, and performed worse than the others. Both models behaved similarly and yielded a consistent transition rate from the egg stage to the larva and pupa stages. This emphasises that the discrete projection matrix and continuous ODE approaches differ slightly, but mostly in propagating different amounts of numerical error during simulation. The use of transition rates, *P* and *G* in Eqns. 4 and 5, merely offers a way to approximate stage transition parameters from life table parameters.

Offering a deterministic model as an explanation to a stochastic system (or a deterministic system with observation error) leaves the noise in observations unaccounted for. This results in a non-zero distance value as the minimum; *i.e.* even the best models achieve a distance no lower than a certain value. Although raising the complexity of a model may result in better fitting, exceedingly complex models risk overfitting, which leads to noise being misinterpreted as signal. Here, no model-parameter combination achieved a distance lower than 331.4.

#### Models offer insights into the system being observed through the distributions of posterior samples

The accuracy of these insights depends heavily on model structure, and thus, the plausibility of its underlying assumptions. We note that, even if the inference is performed with the exact model that generated the output in the first place, a slight deviation from the target should be expected due to the stochasticity of the target model and the finite sample size. The magnitude of this deviation decreases with increasing sample size.

In Table 2, we present the parameter configurations achieving minimum distances in each posterior sample. We observed that the inference resulted in close proximity of the target configuration with the sPop model. The Diff model, on the other hand, implied longer development times for both eggs and larvae. Egg mortality was higher, but larva mortality was lower than expected. The ODE model incorporates instantaneous transition rates, which could roughly translate into exponential decay rates. According to this, the model implied that the expected development time for eggs was 1*/*0.0271 ≈ 37 days and for larvae was 1*/*0.0648 ≈ 15 days. Although egg mortality was close to the expected value, the model implied lower mortality for larvae.

**Table 2.** Best-fitting model-parameter configurations. Models and representative minimum distance parameter values are shown as possible explanations to the observations. The target model and parameter values are given in the final row.

| Model | Parameters (name - value) |  |  |  |  |  |  |  |  |  |  |  |  |
| --- | --- | --- | --- | --- | --- | --- | --- | --- | --- | --- | --- | --- | --- |
| sPop | $p_E$ | 0.0116 | $\mu_{d_E}$ | 25.5 | $\sigma_{d_E}$ | 14.3 | $p_L$ | 0.0547 | $\mu_{d_L}$ | 9.74 | $\sigma_{d_L}$ | 5.19 | |
| Diff | $p_E$ | 0.0244 | $d_E$ | 30.2 | | | $p_L$ | $3.31 \times 10^{-5}$ | $d_L$ | 15.1 | | | |
| ODE | $\beta_E$ | 0.0121 | $\alpha_E$ | 0.0271 | | | $\beta_L$ | 0.0291 | $\alpha_L$ | 0.0648 | | | |
| DDE | $\beta_E$ | 0.0112 | $\alpha_E$ | 5.08 | | | $\beta_L$ | 0.0995 | $\alpha_L$ | 8.04 | | | $\tau_T$ 34.6 |
| sPop (stoch.) | $p_E$ | 0.01 | $\mu_{d_E}$ | 25.0 | $\sigma_{d_E}$ | 15.0 | $p_L$ | 0.05 | $\mu_{d_L}$ | 10.0 | $\sigma_{d_L}$ | 5.0 | |

According to the DDE model, the egg mortality is close to the expected value, however, larva mortality is higher than the target. Interpretation of the development times is less straightforward. The model assumes identical development times for all individuals. When the variable entry modification is not applied, see Proposed model 4: delay differential equations model (DDE), and the model is provided with a development time of, for instance, 25 days, all individuals are incubated for 25 days in the egg stage and transferred to larvae at once. This clearly contradicts the gradual transition seen in the observations. When we applied the variable entry modification, we observed that egg and larva development took approximately 5 and 8 days, respectively, and the observed variation was due to individual eggs commencing development around 35 days apart (Table 2).

The developmental age distribution constructed with variable entry becomes identical for both the egg and larva stages. When this variation is integrated into development times, the model can be interpreted as representing a population of fast and slowly developing individuals. For instance, we observed that a small fraction of the population completed both stages in about 13 days, while a disparate fraction completed development slowly in approximately 48 days. In essence, larva development took about 1.6 times longer than egg development. Consequently, the maximum development times were approximately 18 days for eggs and 30 days for larvae.

### Posterior samples obtained with ABC are weighed equally

When sampling with ABC, any parameter with a distance lower than the threshold is treated equally and assigned the same weight when calculating model predictions [49]. The optimum parameter configuration does not represent the single best fit to the data. The entire posterior should, therefore, be explored and all relevant parameter configurations should be included in predictions. This is often not practical or computationally feasible. Here, we focus on a partial posterior sample, the posterior mode, which is sampled around an optimum value [17].

We plot the posterior samples obtained for the sPop model in Figure 3 together with the marginal frequency distributions and kernel density estimates (KDEs, obtained with the kde function of the ks package in R). The posterior samples for the remaining three models are given in Figures S2 Figure for the Diff model, S3 Figure for the ODE model, and S4 Figure for the DDE model.

**Figure 3.**
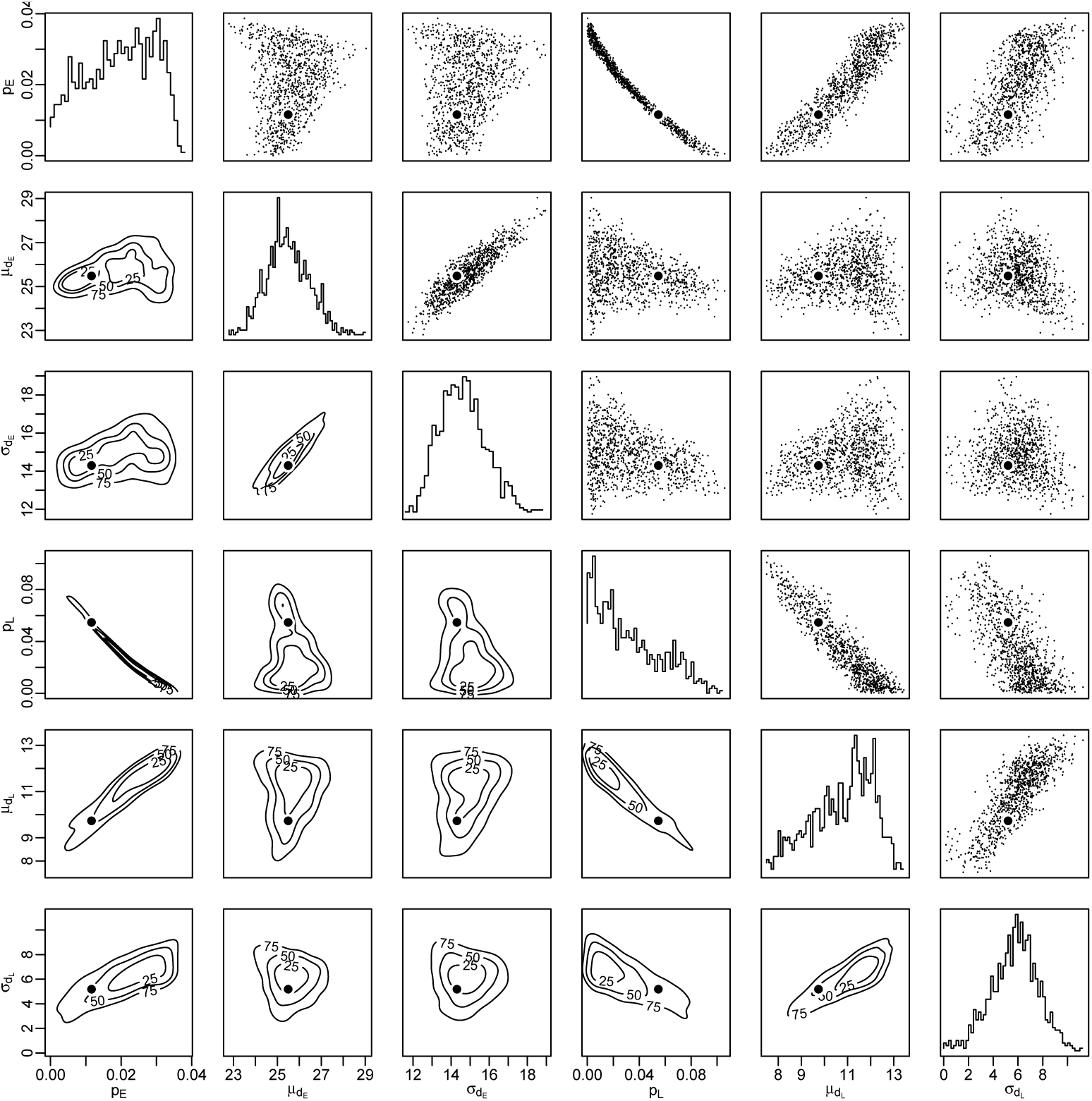
Posterior mode for the sPop model. The figure shows 1000 posterior samples for sPop obtained around the optimum point given in Table 2. Parameter values are shown above, kernel density estimates are plotted below, and marginal frequency distributions are given along the diagonal. The optimum parameter configuration is marked with a solid circle.

A strict negative correlation is evident between the egg and larva mortalities in the posterior samples of the sPop model (Fig. 3). This indicates that the combination of low egg mortality and high larva mortality has the same effect as the opposite configuration. In the vicinity of the target values for mortality, we observed that the samples included all target values for survival and development.

A negative association between the mortalities of egg and larva was also evident in the posterior samples of the DDE model (see S4 Figure). The target values corresponding to the parameterisation of DDE are *β*_*E*_ = 0.01 and *β*_*L*_ = 0.05. We observed that with *ϵ* = 558.181, the samples excluded these values. Similarly, the samples obtained for the ODE and Diff models (with *ϵ* = 1118.7 and *ϵ* = 1041.56, respectively) did not extend over the corresponding target values (*p*_*E*_ = *β*_*E*_ = 0.01 and *p*_*L*_ = *β*_*L*_ = 0.05) with the exception of *β*_*E*_ ranging between 0.008 and 0.014 for the ODE model.

The corresponding target values for development times are *d*_*E*_ = 25 and *d*_*L*_ = 10 for the Diff model and *α*_*E*_ = 0.04 and *α*_*L*_ = 0.1 for the ODE model. We observed that the posterior samples of either model included these values: samples for Diff ranged *d*_*E*_ = 28.28 − 32.30 and *d*_*L*_ = 14.34 − 15.88, and for ODE ranged *α*_*E*_ = 0.024 − 0.032 and *α*_*L*_ = 0.057 − 0.076. We note that the inferred development times were consistently longer than expected when using either the Diff or the ODE models.

Samples for the DDE model implied that egg development time ranges between 2.30 and 6.29 days, while larva development time ranges between 7.32 and 11.26 days. This is accompanied by the variable entry time ranging between 32.94 and 36.23 days. We incorporated variable entry times into development times and focused on total development to facilitate model comparison. According to this, the DDE model implied a median development time in the range of 29.84-31.39 days, however, the target value was 35 days.

### More data are needed to improve the inference

It is possible to further constrain the posterior by adding new observations. For instance, the difference between the total larva and pupa production can be used to pinpoint mortality rates for eggs and larvae.

In order to incorporate new data, we re-configured Eqn. 1 by considering the posterior Pr(*θ* |*δ*) as a prior with regards to new data, *δ*′. Thus,

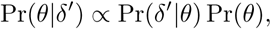

where

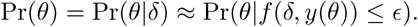

We replaced the likelihood, Pr(*δ*′|*θ*), with a distance function, *f*′, and sampled with an arbitrary threshold, *ϵ*′, as in Eqn. 2.

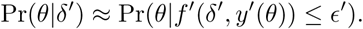

We assigned the new distance function as

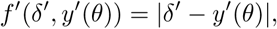

where *y*′(·) is the total number of dead larva predicted by the proposed deterministic model with *θ* and *ϵ*′ = 6.

We observed that for the simulation shown in Figure 1, *δ*′ is, on average, 31 (the values were 25, 31, and 37).

We used the 1000 samples obtained for the sPop model as samples from the prior distribution. We observed that 150 of them yielded dead larvae in the range 25-37, which conforms the distance threshold *ϵ*′ = 6. The samples, as shown in S1 Figure, constrained the posterior distribution for both the egg and larva mortalities around the corresponding target values. The additional datum also constrained the inferred development times to 24.07-26.92 (±12.48-16.43) for eggs and 8.16-11.67 (±0.33-8.47) for larvae.

### Case 2: Inference with an experimental dataset for *Culex quinquefasciatus*

*Culex quinquefasciatus*, commonly known as southern house mosquito, is an important vector of bancroftian filariasis and West Nile virus, and it has been shown capable of transmitting chikungunya and Zika viruses [1, 22, 46]. It is widely distributed across the tropics and sub-tropics.

In 2010, Gunay *et al.* reported a comprehensive study of *Cx. quinquefasciatus* (Say, 1823) [24] at five constant temperatures (15, 20, 23, 27, and 30°C). As described in Gunay *et al.* 2009 and 2010 [23, 24], the study involved regular observations of a total of 750 1^*st*^ instar larvae incubated at 65 ± 5% relative humidity and 14:10 hours of light:dark (L:D) cycle for three consecutive generations in three replicates.

Although the experiment lead to the derivation of several life table parameters, such as adult longevity, generation times, and egg-to-adult survival rates, characteristics of intermediate development stages could not be accurately determined due to overlapping stage durations. Here, we used a subset of these observations (second generation larvae developing at 15°C) to demonstrate the added value of model-based analysis in resolving development and survival characteristics of larvae and pupae (Fig. 4a).

**Figure 4.**
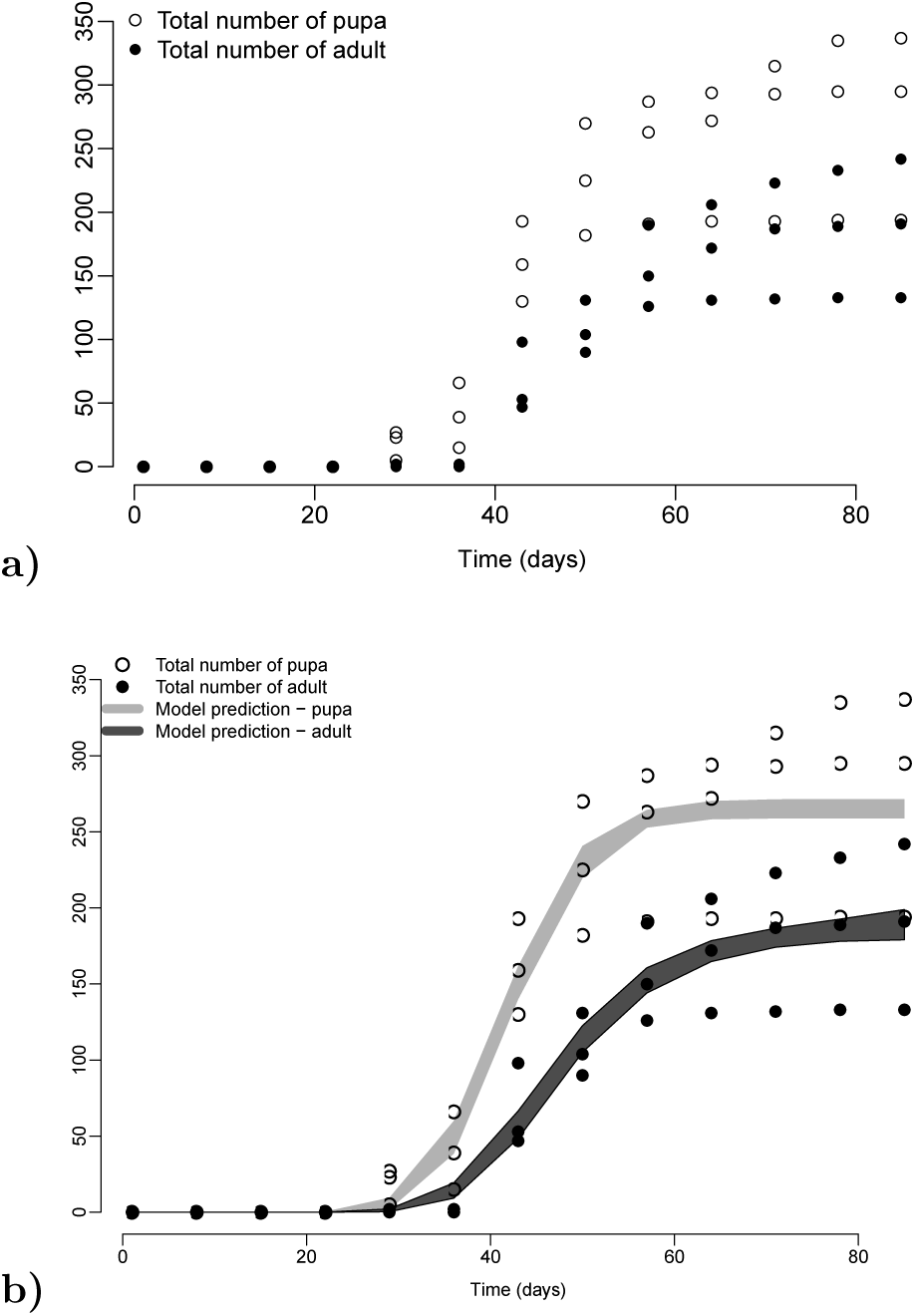
Life table experiment for laboratory-reared *Culex quinquefasciatus*. In (a), the total number of pupa (hollow circles) and adult mosquitoes (solid circles) emerging from 750 larvae are shown in 3 replicates. In (b), the observations are shown with model predictions obtained using the sPop model with 1000 posterior samples. The longitudinal dataset was adopted from Gunay *et al.* (2009) [23].

We developed a deterministic sPop model with two consecutive stages, larvae and pupae, and recorded the number of emerging adults similar to the model in section Proposed model 1: age- and stage-structured difference equations model (sPop). As a result of optimisation and sampling from the optimum posterior mode, we obtained 1000 parameter configurations, which yielded the predictions shown in Figure 4b. The model output seen in the figure corresponds to the expected trajectories of pupae and adults with regard to the three replicates obtained from the laboratory experiments.

We present the posterior samples in Figure 5. A strict positive correlation is evident in the figure between the mean and standard deviation of pupa development time (*σ* = 2.146*µ* − 9.028). With the amount of data utilised, it was not possible to pinpoint pupa development, but, rather, to argue that its standard deviation is roughly twice as much as its mean.

**Figure 5.**
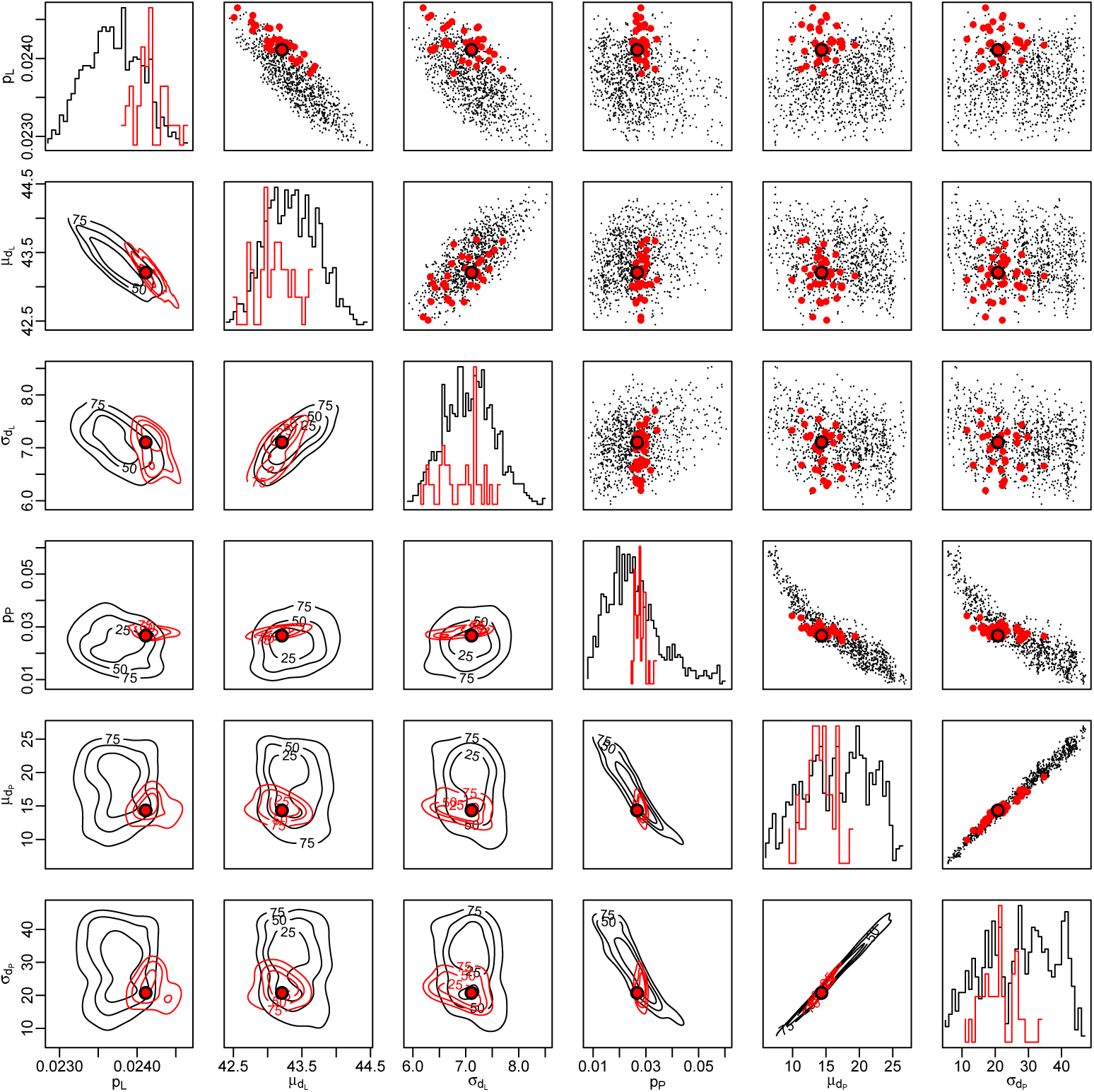
Posterior mode for the sPop model. The figure shows 1000 posterior samples for sPop obtained around the optimum point given in Table 2. The samples yielding larva and pupa mortalities closer than 1% to the observed are marked in red. Parameter values are shown above, kernel density estimates are plotted below, and marginal frequency distributions are given along the diagonal. The optimum parameter configuration is marked with a solid circle.

In addition to the above dataset, Gunay *et al.* reported that on average 255 pupae and 190 adults were produced from 750 larvae at 15°C [23]. By using these observations and following the line of work described in the previous section, we found that 35 of the 1000 posterior samples yielded larva and pupa mortalities closer than 1% to the ones reported (Fig. 5). As a result, we observed that the daily mortality was constrained to 2.42 ± 0.02% for larvae and 2.85 ± 0.21% for pupae.

Gamma distribution with a standard deviation higher than its mean yields a low median. We observed that, although the mean development time of pupa was approximately 14.5 days, the median development time calculated using the inferred parameter values was 5.85 ± 0.68 days. The probability of completing development at 15°C with respect to the time spent for the process are shown in Figure 6 for larva and pupa.

**Figure 6.**
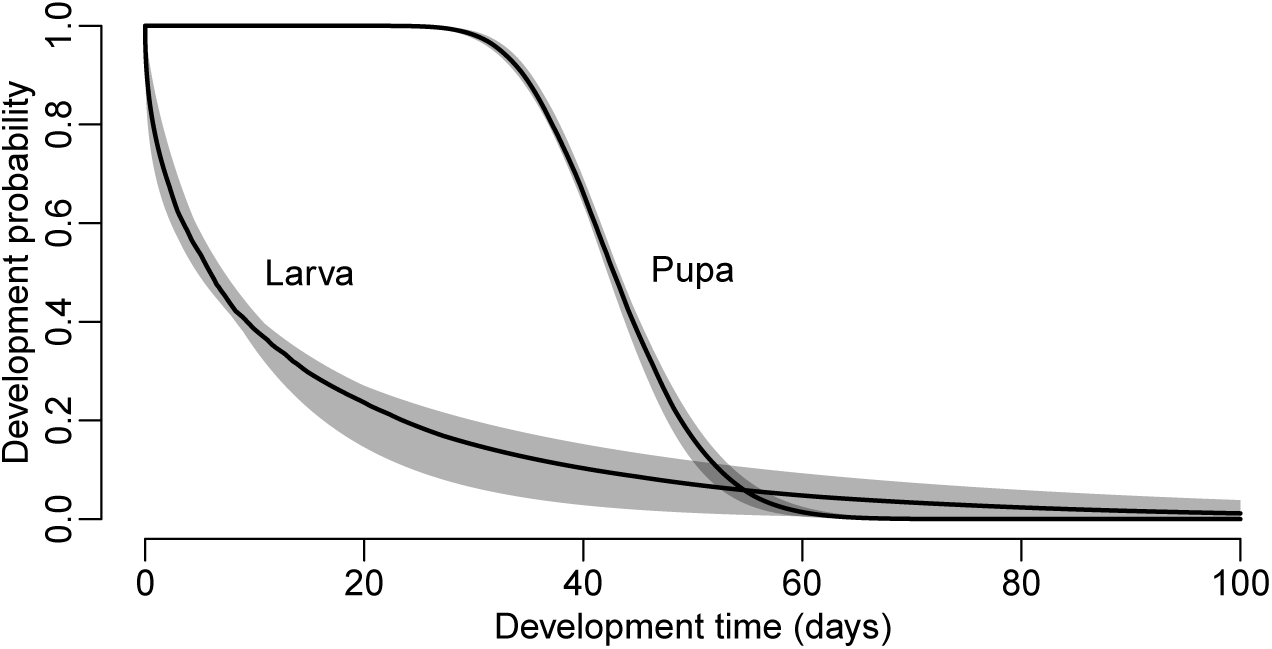
Larva and pupa development at 15°C for *Culex quinquefasciatus* inferred with the sPop model. Development probabilities are estimated using the sPop model and the 35 posterior samples as described in the main text. The medians are given as solid lines and the ranges are given as shades for larvae and pupa.

## Case 3: Inference with an experimental dataset for *Culex pipiens*

In contrast to its southern neighbour, *Culex pipiens* is known as the northern house mosquito as it is widely distributed across the temperate countries in North America, Europe, Asia, and North and East Africa [9, 18]. It is an important vector of West Nile and St. Louis encephalitis viruses and filarial worms [18].

In 2019, a life table experiment was conducted by Petrić’s group with a laboratory-reared population of *Culex pipiens* biotype molestus to examine its environmental dependence. Four batches of eggs (180, 177, 161, and 153 eggs) were prepared and their development was observed at room temperature. Each batch was kept in 1.5 L water at 19.9 ± 1.0°C and 38.8 ± 2.5% relative humidity. Following hatching and larva development, pupa production was recorded each day until all larvae either died or developed. We present the observations in Figure 7(a).

**Figure 7.**
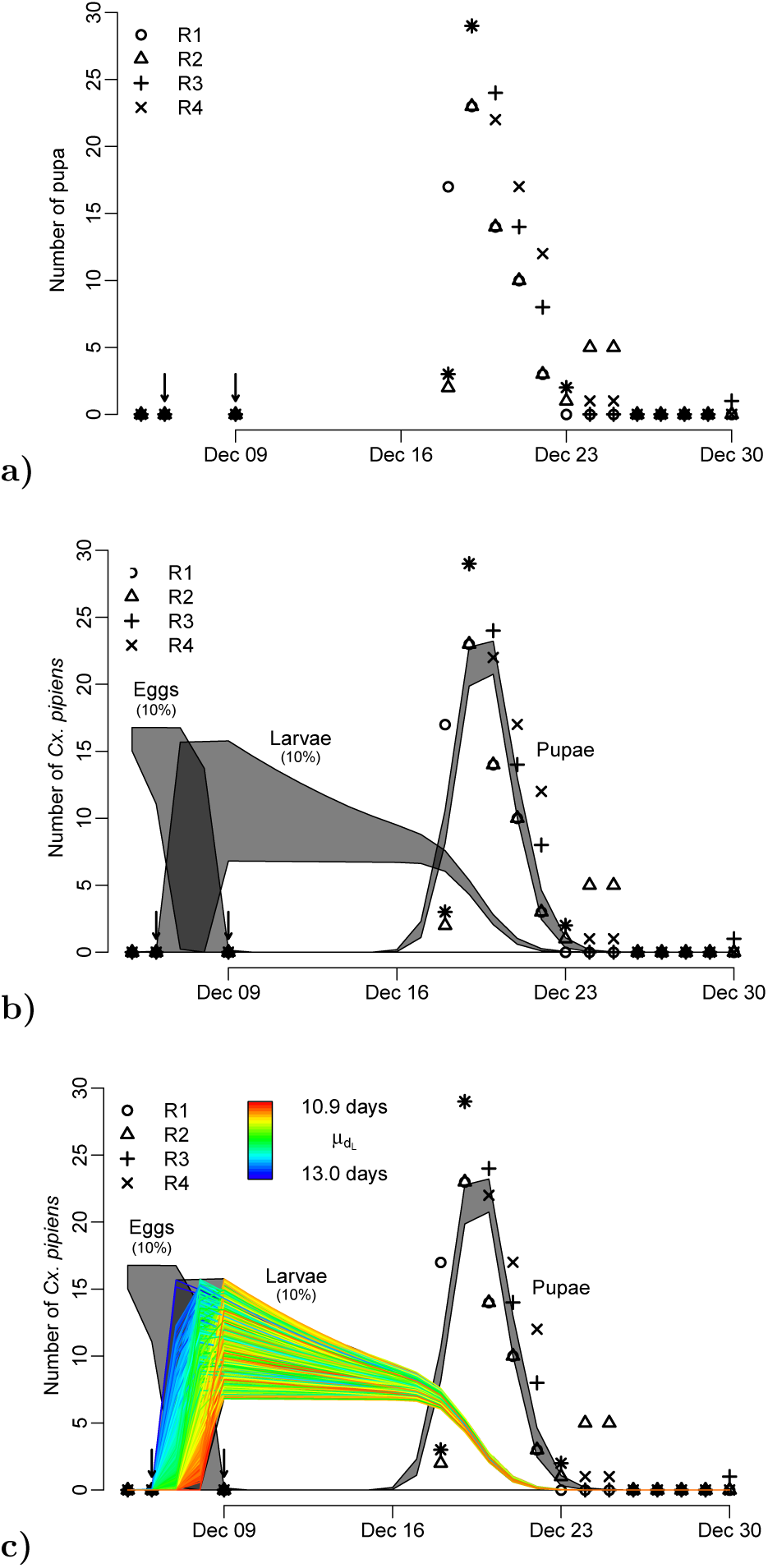
Life table experiment for laboratory-reared *Culex pipiens* at room temperature. In (a), the total number of pupa emerging from the four batches of egg are shown. In (b), the observations are shown with model predictions (minimum and maximum pupa production per day) obtained using the sPop model with 1000 posterior samples. In (c), model outputs for larvae are plotted and marked for a range of 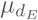 values from the posterior.

While pupa production was recorded, counts of viable eggs or larvae were not. Instead, the group noted that none of the eggs hatched on the second day and all the eggs hatched after four days. We marked the times of these two additional observations with arrows in the figure.

In order to represent this experiment, we developed a deterministic two-stage sPop model with egg and larva stages where pupa production is recorded daily. By default, sPop is a daily time-step difference equations model. However, due to the rapid hatching of the eggs, we employed a time unit of quarter days where each iteration accounts for 6 hours rather than 1 day. Following each simulation, we calculated the daily mean number of viable eggs and larvae and the total pupa production for each day.

As the distance function, we employed the sum of squares distance for pupa production and included the two additional observations, *i.e.*

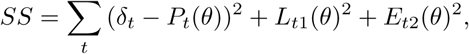

where *δ*_*t*_ refers to the number of pupa produced on day *t, P*_*t*_(*θ*) is the corresponding model simulation with parameter *θ, L*_*t*1_(*θ*) is the simulated number of larvae on *t*1 (1 day after oviposition), and *E*_*t*2_(*θ*) is the simulated number of eggs on *t*2 (4 days after oviposition).

Following the same optimisation and posterior sampling procedure as in previous sections, we observed that the model closely represented the bulk of observations on pupa production (Fig. 7(b)). Due to the lack of detailed records on earlier stages, a wide range of uncertainty was assigned for egg and larva development.

As an added value, the model offers a mean to drastically improve precision. A key observation is revealed upon close examination of the posterior mode. As seen in Figure 7(c), the first three days beginning with the observation of first egg hatching seem to offer the ability to resolve larva development time. If the mean development time 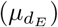 is around 11 days, the peak in the number of larva realises later than if 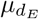 was around 13 days.

For instance, let us assume that on December 7^*th*^, the first day of egg hatching, the initial number of larvae were counted and found to be between 15 and 35 in each of the 4 replicates. This observation constrains both egg and larvae mean development times around 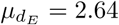 and 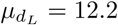 days, respectively while not significantly affecting the parameters for survival and standard deviation (*p*_*E*_, *p*_*L*_, 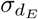, and 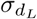).

**Figure 8.**
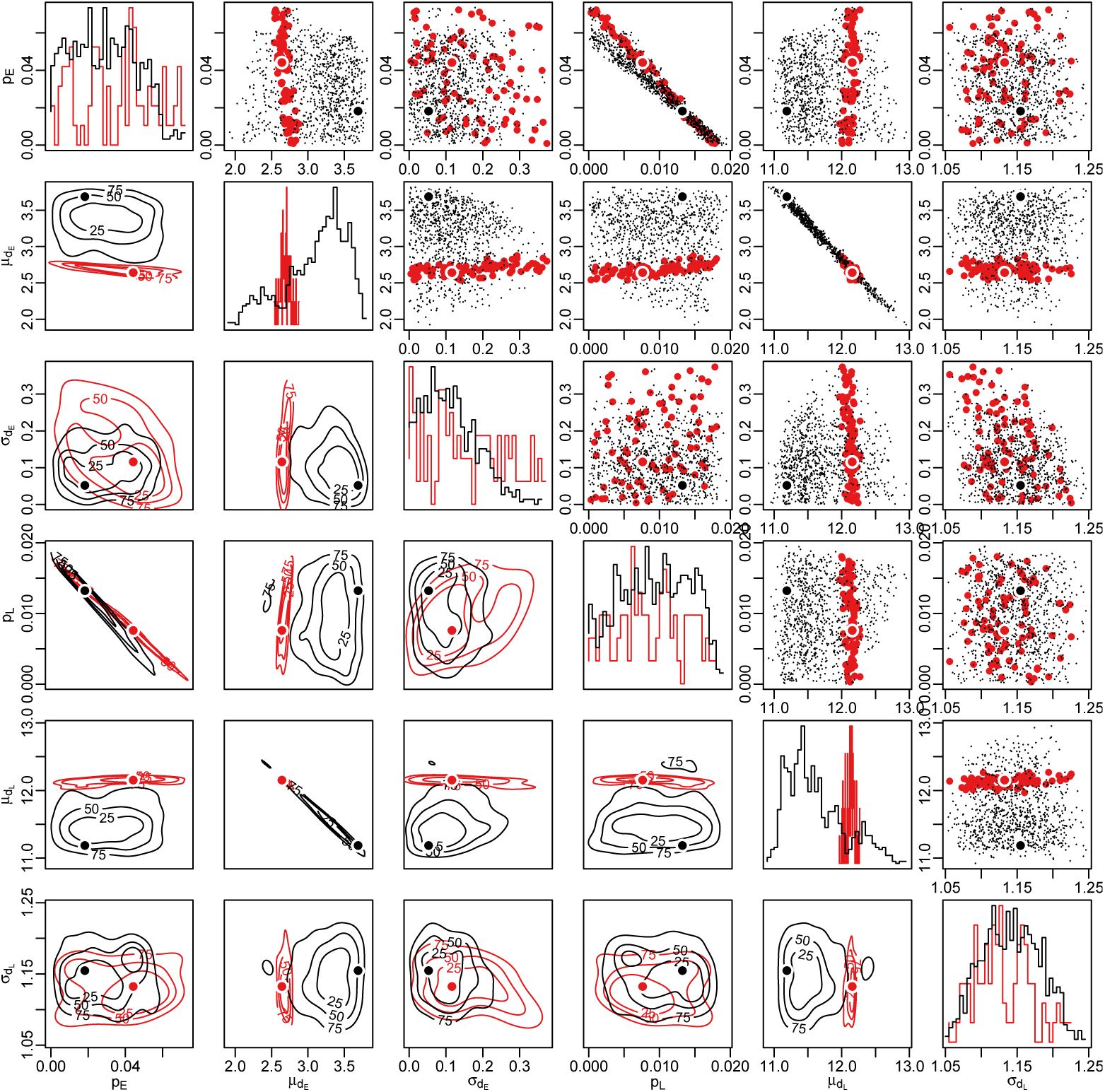
Posterior mode for the sPop model of *Culex pipiens*. The figure shows 1000 posterior samples, 88 of which are marked in red and correspond to a larva count of 15-35 on December 7^*th*^. Parameter values are shown above, kernel density estimates are plotted below, and marginal frequency distributions are given along the diagonal. The optimum parameter configuration is marked with two solid circles for each group.

## Discussion

This study demonstrates that age- and stage-structured population dynamics modelling can help estimate developmental characteristics of intermediate life stages from laboratory experiments. With this approach, we amplify the amount of information discernable from canonical experimental designs and suggest ways to improve accuracy.

Insect development is a complex cycle, which requires simplifying assumptions in incorporating the level of this complexity in mathematical models. In order to gain meaningful insights, any model proposed should align well with the underlying biological processes governing development. We demonstrated that the inclusion of population heterogeneity in the form of age structure leads to time delays and suffices to represent the process of development.

In contrast, inappropriate models may lead to misleading conclusions. We evaluated a representative set of frequently encountered modelling tools against the best case scenario, where the system which produced the data is fully known and there is no unaccounted external noise in the observations. Consequently, none of these tools, except the sPop model, represented the development process in required detail.

Using the sPop model, we demonstrated that observations during stage transitions offer more information on immature stage development than the ones during the steady progression of a particular stage. Intrinsic noise contaminates both phases, but often outweighs the signal during stage progression. Focusing observations on key time points could provide the necessary information especially in limited resource cases.

Two types of observations can help isolate and focus on intermediate stage development: The number of viable individuals and the total production of a particular stage. The former informs about the rate of development and survival while the latter helps to interrupt the time series and investigate earlier stages in isolation. Despite its advantages, variability in the records of total production accumulates in time and reaches the maximum at the end. Although this may not be evident in the time series of a single experiment, variability is exposed when multiple biological replicates are compared. This introduces bias to the inference especially when the variability is dismissed with the use of a generic distance function (such as the sum of squares).

We suggest that further optimisations of this procedure should consider revising the distance measure to better reflect the underlying statistical distribution at each time point and to accommodate any anticipated time correlation. We suggest that designing multiple biological replicates and increasing the frequency of observations during key time intervals would significantly increase the information content of the data.

The proposed approach is readily applicable for complex experimental designs, such as variable-temperature life table experiments, which had to be reduced and simplified previously. The approach also allows for developing population dynamics models primarily informed by field observations. As shown previously [16, 17], such models can provide valuable predictive ability under field conditions across a broad spatial range.

## Conclusion

Life tables have long been used to understand population dynamics and environmental dependence [10]. However, their resolution is limited given overlapping development stages, population heterogeneity, and the intricate interactions between individuals and their environment. Laboratory experiments under controlled conditions often risk disrupting such interactions, influencing natural behaviour, and affecting the outcome. We demonstrated that model-based inference can be a valuable tool in estimating life characteristics from complex and, often, incomplete observations while minimising the requirement of disruptive interventions.

We conclude that an appropriate model structure is necessary to derive valid conclusions. Model structure should be informed by valid assumptions arising from prior knowledge, and used to complement longitudinal observations. With its ability to follow age-structure within stage-structured populations, the sPop model represents heterogeneity and time delay resulting from development, and offers a viable alternative to the mainstream approaches.

Parameter values inferred from life table experiments using the sPop model are readily applicable as prior knowledge in further iterations of population dynamics models. The resulting models can help to understand underlying biological processes including environmental dependence and disease transmission, and be used in projecting large-scale trends in climate-driven habitat suitability and disease risk.

## Acknowledgments

We thank Sean Wu and Jacob Mendel for valuable discussions and critical comments on the earlier version of this manuscript. We also thank Asena Çapkın and İrem Kaşıkçı for helpful discussions.

This publication has been produced within the framework of the EMME-CARE project which has received funding from the European Union’s Horizon 2020 Research and Innovation Programme, under Grant Agreement No. 856612 and the Cyprus Government. The sole responsibility of this publication lies with the authors. The European Union is not responsible for any use that may be made of the information contained therein.

The work was done within the framework of AIM-COST Action CA17108 “*Aedes* Invasive Mosquitoes”.

The work was partially supported by The Armed Forces Health Surveillance Center, Global Emerging Infections Surveillance and Response System (AFHSC-GEIS) (W81XWH-18-C-0190-P0001), United States.

KE acknowledges the Cy-Tera Project (NEW INFRASTRUCTURE/STRATEGIC/0308/31), which is co-funded by the European Regional Development Fund and the Republic of Cyprus through the Research Promotion Foundation, for providing the computational resources used in this research.

The funders had no role in study design, data collection and analysis, decision to publish, or preparation of the manuscript.

## Supporting Information

### S1 Text

#### The Metropolis-Hastings Markov chain Monte Carlo (MCMC) algorithm (R script)

The following code was inspired by the work of Darren Wilkinson (https://darrenjw.wordpress.com/2010/08/15/metropolis-hastings-mcmc-algorithms/).

~~~
mcmc <- function (pr, fun, lower, upper, kernel.sd,
      niter =1000, thin =10, sig =1 .0, verbose= FALSE) {
 acc <- 0
 vec <- NULL
 prn <- pr
 scrn <- scr <- fun (pr)
 for (n in 1: niter) {
   repeat {
    prn <- rnorm (length (pr), mean = pr, sd= kernel.sd)
    if (all(prn >= lower) && all(prn <= upper)) break
   }
   scrn <- fun (prn)
   if (! is.nan (scrn) && (scrn < scr ‖ log (runif (1)) < (scr - scrn)/ sig)) {
    pr <- prn
    scr <- scrn
    acc <- acc + 1
   }
  vecn <- c(scr, pr)
  if (n %% thin == 0) {
    vec <- rbind (vec, vecn)
    if (verbose) cat(sprintf(“% d, % g, % s\ n”,n, acc, paste(vecn, collapse=“,”)))
    acc <- 0
  }
 }
 colnames(vec)<-c(“score”, sprintf(“X% d”, 1: length (pr)))
 return (data.frame (vec, row.names= NULL))
}
~~~

### S2 Text

#### The approximate Bayesian computation - sequential Monte Carlo (ABC-SMC) algorithm (R script)

The following code was inspired by the works of Toni *et al.* (2008) and Liepe *et al.* (2010) [33, 49].

~~~
abc.smc <- function (pr, score, eps, size, lower, upper, kernel.sd, niter, verbose= TRUE) {
  scr <- score(pr)
  mat <- matrix (rep (c(1 / size, 0, scr, pr), size), nrow = size, byrow = TRUE)
  mat.new <- matrix (NA, nrow = nrow (mat), ncol= ncol(mat))
  for (i in 1: niter) {
   for (n in 1: nrow (mat.new)) {
      fcount <- 0
      repeat {
       v <- mat[ sample (1: nrow (mat),1, prob = mat[,1], replace= TRUE),]
       pr.new <- rnorm (length (v)-3, mean = v [ -(1:3)], sd= kernel.sd)
       scr <- score(pr.new)
       if (! is.nan (scr) && (scr < eps) &&
          all(pr.new >= lower) && all(pr.new <= upper)) break
       fcount <- fcount + 1
      }
      if (verbose) print(c(i,n, fcount))
      w <- 1 / sum (apply (mat, 1, function (v)
        v [1]* exp (sum (dnorm (pr.new, mean = v [ -(1:3)], sd= kernel.sd, log = TRUE)))))
      mat.new [n,] <- c(w, fcount, scr, pr.new)
   }
   mat.new [,1] <- mat.new [,1] / sum (mat.new [,1])
   mat <- mat.new
   if (verbose) {
      cat(paste(apply (mat,1, paste, collapse=“,”), collapse=“\ n”), sep =“\ n\ n”)
   }
  }
  return (mat)
}
~~~

### S3 Text

#### The stochastic sPop model (R script)

The model was used to simulate hypothetical experimental data of a given repetitions. The output of the function is a data.frame of weekly model output starting from day 1, the day after initiation.

~~~
fun.sPop.st <- function (pr, rpt =3) {
  require(albopictus)
  ret <- NULL
  for (r in 1: rpt) {
    print(r)
    egg <- spop (stochastic = TRUE, prob = ‘gamma’)
    larva <- spop (stochastic = TRUE, prob = ‘gamma’)
    add (egg) <- data.frame (number = 100)
    larvad <- 0
    pupa <- 0
    for (n in 1:98) {
       iterate(egg) <- data.frame (death = pr[1], dev _ mean = pr[2], dev _ sd= pr [3])
       iterate(larva) <- data.frame (death = pr[4], dev _ mean = pr[5], dev _ sd= pr [6])
       larvad <- larvad + dead (larva)
       pupa <- pupa + developed (larva)
       add (larva) <- data.frame (number= developed (egg))
       if (n%% 7== 1) {
         ret <- rbind (ret, c(r,n, size(larva), larvad, pupa))
       }
     }
  }
  colnames(ret) <- c(“rep “,” day “,” larva”,” larvad “,” pupa”)
  ret <- data.frame (ret)
  return (ret)
}
~~~

### S4 Text

#### The deterministic sPop model (R script)

The output of the function is a data.frame of weekly model output starting from day 1, the day after initiation. The output comprise the following in the given order:

day Day of observation (weekly)

stage2 Viable number of individuals in stage 2 at the time of observation cum.stage2 Total number of individuals in stage 2 at the time of observation

cum.stage2.dead Total number of non-viable individuals in stage 2 at the time of observation cum.stage3 Total number of individuals in stage 3 at the time of observation

~~~
fun.sPop <- function (pr, tm, init) {
  require(albopictus)
  stage1 <- spop (stochastic = FALSE, prob = ‘gamma’)
  stage2 <- spop (stochastic = FALSE, prob = ‘gamma’) cum.stage2 <- 0
  cum.stage2 .dead <- 0
  cum.stage3 <- 0
  add (stage1) <- data.frame (number= init)
  ret <- NULL
  for (n in 1: max (tm)) {
    iterate(stage1) <- data.frame (death = pr[1], dev _ mean = pr[2], dev _ sd= pr [3])
    iterate(stage2) <- data.frame (death = pr[4], dev _ mean = pr[5], dev _ sd= pr [6])
    new.stage2 <- developed (stage1)
    new.stage3 <- developed (stage2)
    dead.stage2 <- dead (stage2)
    add (stage2) <- data.frame (number= new.stage2)
    cum.stage2 <- cum.stage2 + new.stage2
    cum.stage2 .dead <- cum.stage2 .dead + dead.stage2
    cum.stage3 <- cum.stage3 + new.stage3
    ret <- rbind (ret, c(n, size(stage2), cum.stage2, cum.stage2.dead, cum.stage3))
  }
  colnames(ret) <- c(“day “,” stage2 “,” cum.stage2 “,” cum.stage2 .dead “,” cum.stage3 “)
  ret <- data.frame (ret)
  ret <- ret[ ret$ day % in% tm,]
  return (ret)
}
~~~

### S5 Text

#### The deterministic Diff model (R script)

The output of the function is a data.frame of weekly model output starting from day 1, the day after initiation. The output comprise the following in the given order:

day Day of observation (weekly)

stage2 Viable number of individuals in stage 2 at the time of observation

cum.stage2 Total number of individuals in stage 2 at the time of observation

cum.stage3 Total number of individuals in stage 3 at the time of observation

~~~
fun.Diff <- function (pr, tm, init) {
  stage1 <- init
  stage2 <- 0
  cum.stage2 <- 0
  cum.stage3 <- 0
  ret <- NULL
  p1 <- 1 - pr [1]
  p3 <- 1 - pr [3]
  pd <- p1 ^ pr [2]
  PE <- (p1 - pd)/(1 - pd)
  pl <- p3 ^ pr [4]
  PL <- (p3 - pl)/(1 - pl)
  GE <- (1 - p1)* pd/(1 - pd)
  GL <- (1 - p3)* pl/(1 - pl)
  for (n in 1: max (tm)) {
    new.stage2 <- GE * stage1
    new.stage3 <- GL * stage2
    cum.stage3 <- cum.stage3 + new.stage3
    cum.stage2 <- cum.stage2 + new.stage2
    stage2 <- new.stage2 + PL * stage2
    stage1 <- PE * stage1
    if (n % in% tm) {
      ret <- rbind (ret, c(n, stage2, cum.stage2, cum.stage3))
    }
  }
  colnames(ret) <- c(“day “,” stage2 “,” cum.stage2 “,” cum.stage3 “)
  ret <- data.frame (ret)
  return (ret)
}
~~~

### S6 Text

#### The deterministic ODE model (R script)

The output of the function is a data.frame of weekly model output starting from day 1, the day after initiation. The output comprise the following in the given order:

day Day of observation (weekly)

stage2 Viable number of individuals in stage 2 at the time of observation

cum.stage2 Total number of individuals in stage 2 at the time of observation

cum.stage3 Total number of individuals in stage 3 at the time of observation

~~~
fun.ODE <- function (pr, tm, init) {
  require(deSolve)
  fun Det <- function (t, y, pr) {
    new.stage2 <- pr [2] * y [1]
    new.stage3 <- pr [4] * y [2]
    d Stage1 <- - new.stage2 - pr [1] * y [1]
    d Stage2 <- new.stage2 - new.stage3 - pr [3] * y [2]
    d Cum.stage2 <- new.stage2
    d Cum.stage3 <- new.stage3
    return (list(c(dStage1, dStage2, dCum.stage2, d Cum.stage3)))
  }
  ret <- data.frame (ode(y = c(init, 0, 0, 0),
        times = tm,
        func = funDet,
        parms = pr)[, c(1, 3, 4, 5)])
  colnames(ret) <- c(“day “,” stage2 “,” cum.stage2 “,” cum.stage3 “)
  return (ret)
}
~~~

### S7 Text

#### The deterministic DDE model (R script)

The output of the function is a data.frame of weekly model output starting from day 1, the day after initiation. The output comprise the following in the given order:

day Day of observation (weekly)

stage2 Viable number of individuals in stage 2 at the time of observation

cum.stage2 Total number of individuals in stage 2 at the time of observation

cum.stage3 Total number of individuals in stage 3 at the time of observation

~~~
fun.DDE <- function (pr, tm, init) {
  require(stagePop)
  ccFunctions <- list(
    death Func= function (stage, x, time, species, strain){
      if (species ==1) return (c(pr [1], 0)[ stage ])
      return (c(pr[1], pr [3], 0)[ stage ])
    },
    duration Func = function (stage, x, time, species, strain){
      if (species ==1) return (c(pr[2], Inf)[ stage ])
      return (c(pr[2], pr[4], Inf)[ stage ])
    },
    repro Func= function (x, time, species, strain){ return (0)},
    emigration Func = function (stage, x, time, species, strain){ return (0)},
    immigration Func = function (stage, x, time, species, strain){
       v <- 0
      if (stage ==1 && pr[5] >0 && time >=0 && time <= pr [5]) {
         v <- init/ pr [5]
      }
      return (v)
    }
 )
  out <- pop Model(
    num Species = 2,
    num Stages = c(2, 3),
    timeDepend Loss = c(TRUE, TRUE),
    timeDepend Duration = c(FALSE, FALSE),
    ICs = list(matrix (0, ncol=1, nrow =2), matrix (0, ncol=1, nrow =3)),
    timeVec = seq (0, max (tm),0 .1),
    rateFunctions = ccFunctions,
    stageNames = list(c(“Stage1 “,” Cum.stage2 “),
            c(“Stage1 “,” Stage2 “,” Cum.stage3 “)),
    speciesNames = c(“Vector1 “,” Vector2 “),
    saveFig = FALSE,
    plotFigs = FALSE
 )
  ret <- data.frame (out[ out [,1] % in% tm,
           c(“time”,
           “Vector2 .Stage2 “,
           “Vector1 .Cum.stage2 “,
           “Vector2 .Cum.stage3 “)])
  colnames(ret) <- c(“day “,” stage2 “,” cum.stage2 “,” cum.stage3 “)
  return (ret)
}
~~~

---

**S1 Figure.**
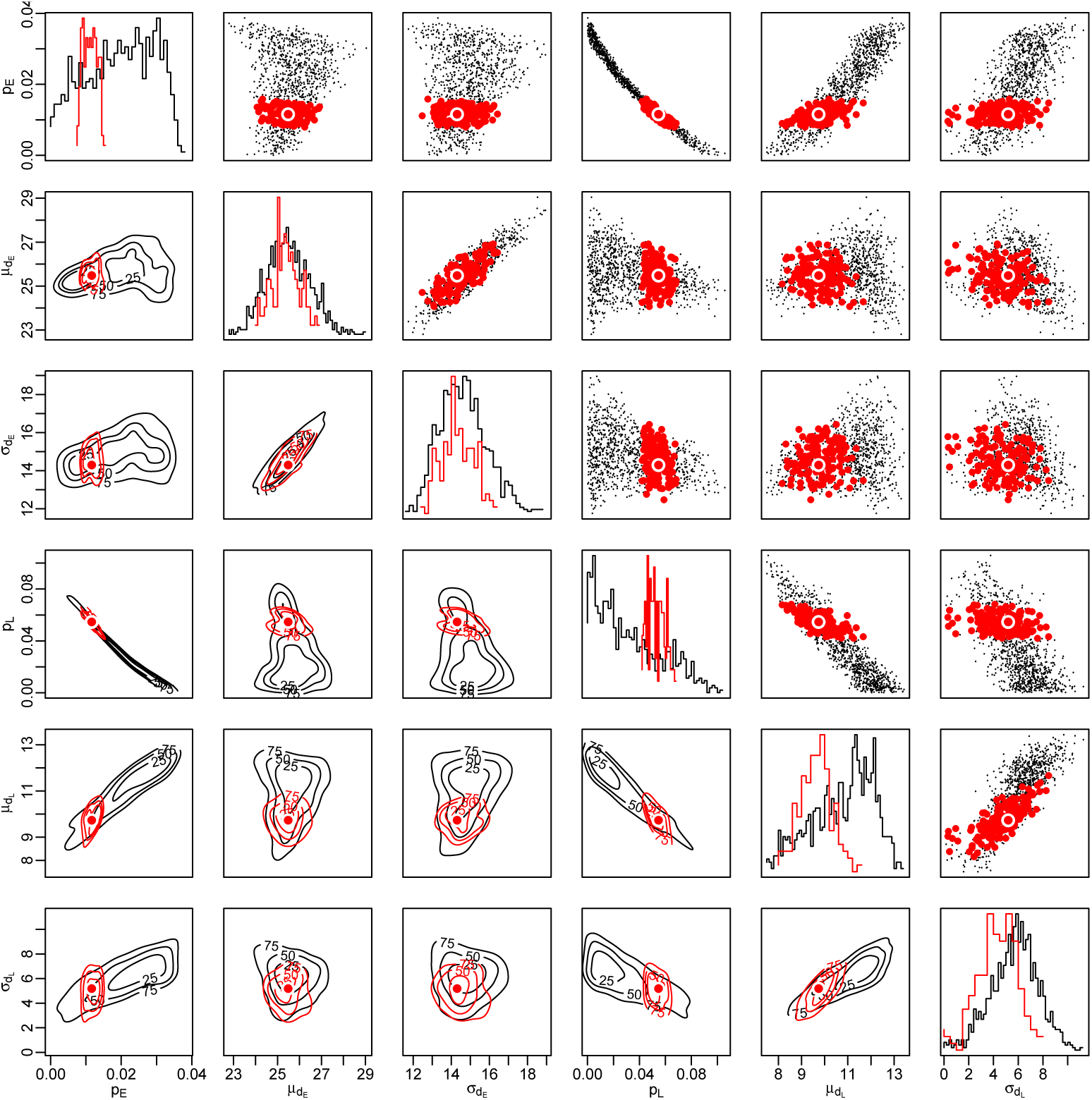
The posterior samples for the sPop model. The 1000 parameter configurations inferred without using the number of dead larvae are plotted in black, and the 150 of them inferred using the number of dead larvae are plotted in red. The parameter configurations are plotted in pairs in the figures above the diagonal. Below the diagonal, kernel density estimates are given. Marginal frequency distributions for each parameter are given along the diagonal. The parameter configuration with the minimum distance is marked with a solid circle.

**S2 Figure.**
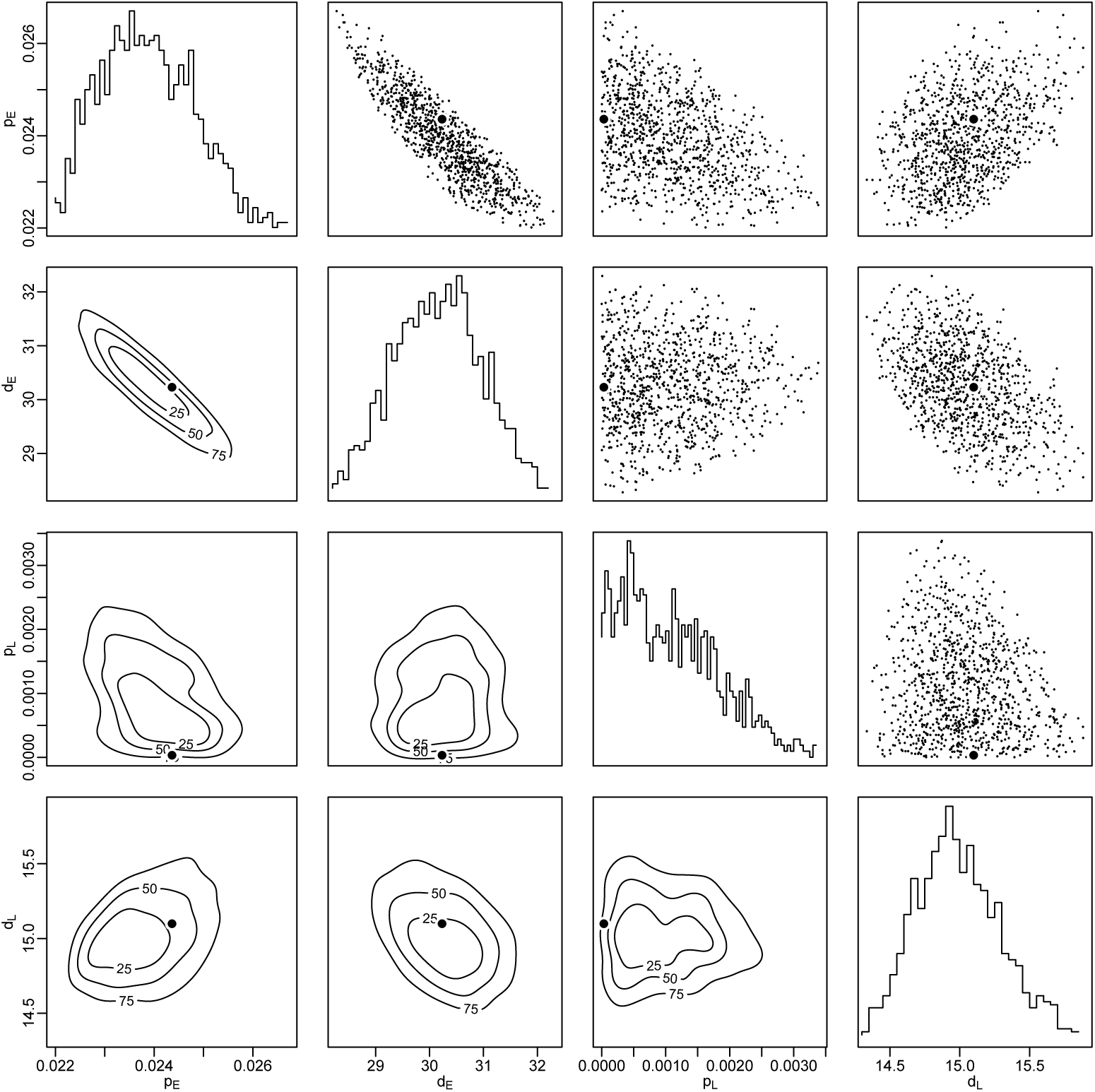
The posterior samples for the Diff model. 1000 parameter configurations are plotted in pairs in the figures above the diagonal. Below the diagonal, kernel density estimates are given. Marginal frequency distributions for each parameter are given along the diagonal. The parameter configuration with the minimum distance is marked with a solid circle.

**S3 Figure.**
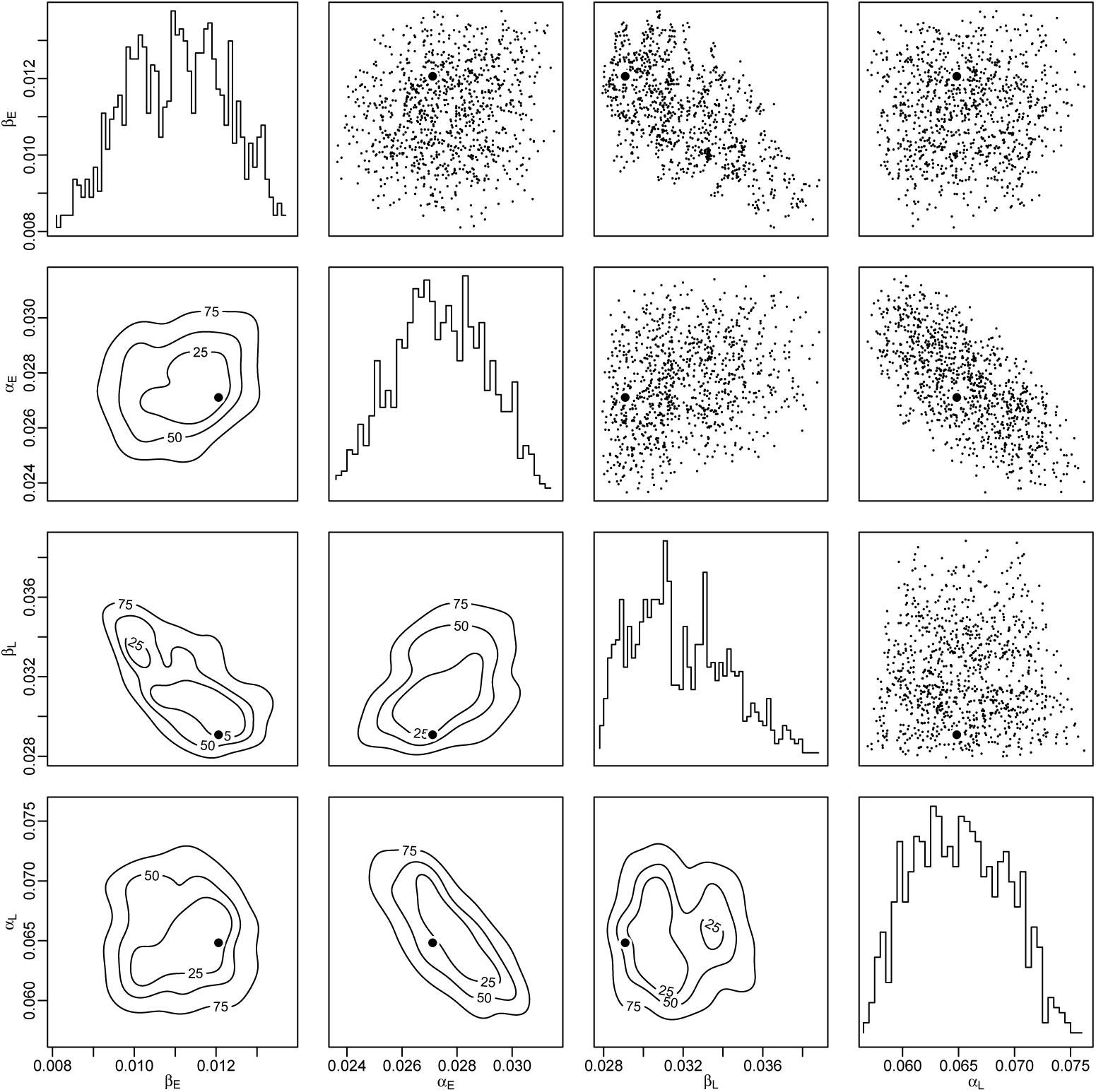
The posterior samples for the ODE model. 1000 parameter configurations are plotted in pairs in the figures above the diagonal. Below the diagonal, kernel density estimates are given. Marginal frequency distributions for each parameter are given along the diagonal. The parameter configuration with the minimum distance is marked with a solid circle.

**S4 Figure.**
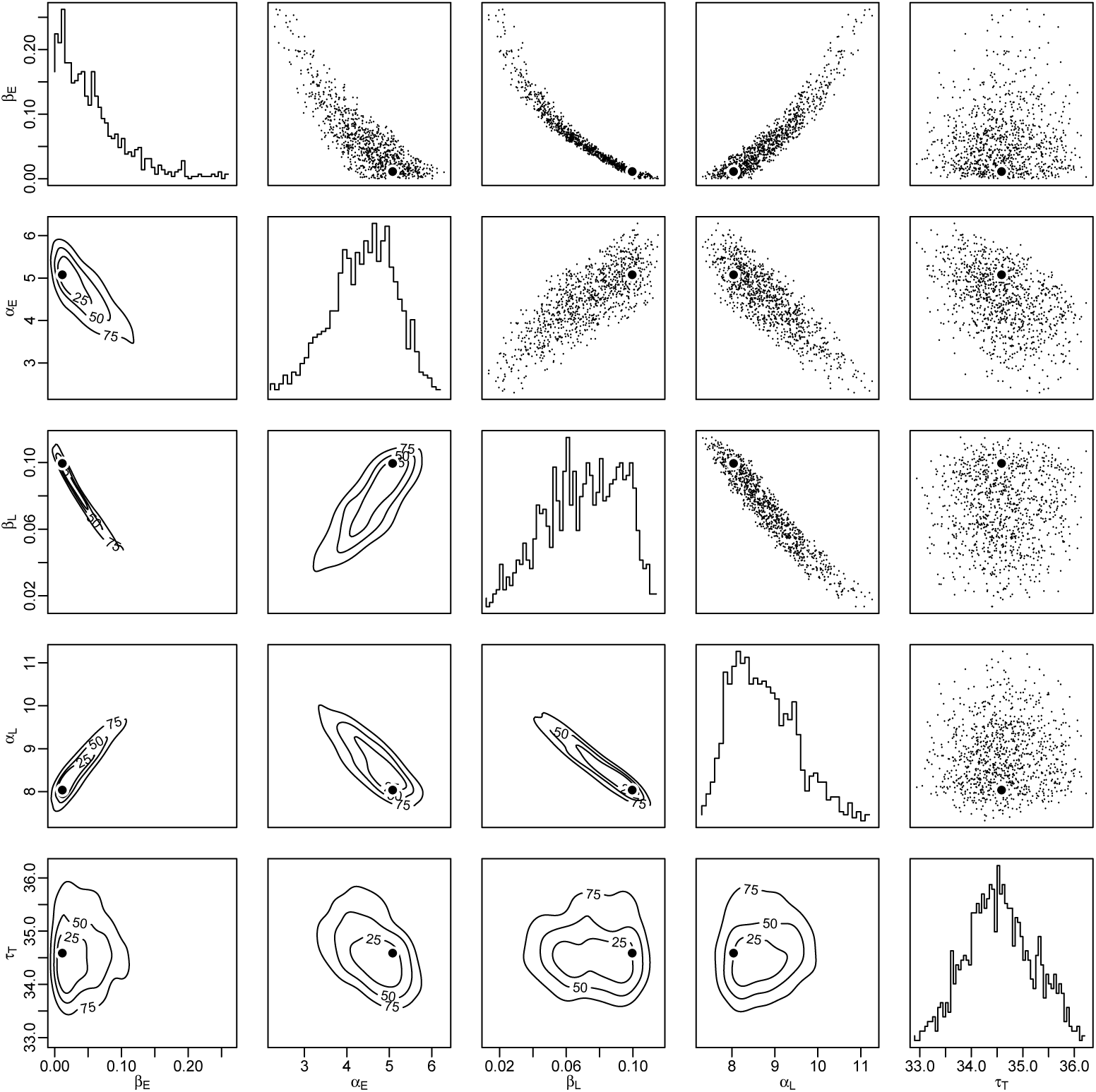
The posterior samples for the DDE model. 1000 parameter configurations are plotted in pairs in the figures above the diagonal. Below the diagonal, kernel density estimates are given. Marginal frequency distributions for each parameter are given along the diagonal. The parameter configuration with the minimum distance is marked with a solid circle.

## References

1. N. Becker, D. Petric, M. Zgomba, C. Boase, M. Madon, C. Dahl, and A. Kaiser. Mosquitoes and Their Control, volume 57. 2010.

2. A. Belen and B. Alten. Variation in life table characteristics among Populations of Phlebotomus papatasi at different altitudes. Journal of Vector Ecology, 31(1):35–44, jun 2006.

3. T. Bellows and R. Van Driesche. Life Table Construction and Analysis for Evaluating Biological Control Agents. In Handbook of Biological Control, pages 199–223. Academic Press, 1999.

4. M. Q. Benedict. Methods in Anopheles Research. MR4 - BEI Resources, 2014.

5. F. Brauer and C. Castillo-Chavez. Mathematical models in population biology and epidemiology. Springer, 2 edition, 2010.

6. S. Brooks, A. Gelman, G. L. Jones, and X. L. Meng. Handbook of Markov Chain Monte Carlo. 2011.

7. B. Calderhead. A general construction for parallelizing Metropolis-Hastings algorithms. Proceedings of the National Academy of Sciences of the United States of America, 111(49):17408–17413, dec 2014.

8. C. Christiansen-Jucht, K. Erguler, C. Y. Shek, M.-G. Basáñez, and P. E. Parham. Modelling anopheles gambiae s.s. population dynamics with temperature- and age-dependent survival. International Journal of Environmental Research and Public Health, 12(6):5975–6005, 2015.

9. A. T. Ciota and L. D. Kramer. Vector-virus interactions and transmission dynamics of West Nile virus. Viruses, 5(12):3021–3047, 2013.

10. D. Cox. Regression models and life tables. J. R. Stat. Soc. B, 34(2):187–220, 1972.

11. D. T. Crouse, L. B. Crowder, and H. Caswell. A stage-based population model for loggerhead sea turtles and implications for conservation. Ecology, 68(5):1412–1423, 1987.

12. H. Delatte, G. Gimonneau, A. Triboire, and D. Fontenille. Influence of temperature on immature development, survival, longevity, fecundity, and gonotrophic cycles of Aedes albopictus, vector of chikungunya and dengue in the Indian Ocean. Journal of Medical Entomology, 46(1):33–41, jan 2009.

13. Edda Klipp, W. Liebermeister, C. Wierling, and A. Kowald. Systems Biology. Wiley-VCH Verlag GmbH & Co. KGaA, 2016.

14. K. Erguler. sPop: Age-structured discrete-time population dynamics model in C, Python, and R. F1000Research, 7:1220, 2018.

15. K. Erguler, N. N. L. Chandra, Y. Proestos, J. Lelieveld, G. G. K. Christophides, and P. P. E. Parham. A large-scale stochastic spatiotemporal model for Aedes albopictus-borne chikungunya epidemiology. Plos One, 12(3):e0174293, 2017.

16. K. Erguler, I. Pontiki, G. Zittis, Y. Proestos, V. Christodoulou, N. Tsirigotakis, M. Antoniou, O. E. Kasap, B. Alten, and J. Lelieveld. A climate-driven and field data-assimilated population dynamics model of sand flies. Scientific Reports, 9(1):2469, dec 2019.

17. K. Erguler, S. E. Smith-Unna, J. Waldock, Y. Proestos, G. K. Christophides, J. Lelieveld, and P. E. Parham. Large-scale modelling of the environmentally-driven population dynamics of temperate aedes albopictus (Skuse). PLoS ONE, 11(2), 2016.

18. A. Farajollahi, D. M. Fonseca, L. D. Kramer, and A. Marm Kilpatrick. “Bird biting” mosquitoes and human disease: A review of the role of Culex pipiens complex mosquitoes in epidemiology. Infection, Genetics and Evolution, 11(7):1577–1585, 2011.

19. L. C. Farnesi, H. C. Vargas, D. Valle, and G. L. Rezende. Darker eggs of mosquitoes resist more to dry conditions: Melanin enhances serosal cuticle contribution in egg resistance to desiccation in Aedes, Anopheles and Culex vectors. PLoS Neglected Tropical Diseases, 11(10):1–20, 2017.

20. L. Gavotte, D. R. Mercer, R. Vandyke, J. W. Mains, and S. L. Dobson. Wolbachia infection and resource competition effects on immature Aedes albopictus (Diptera: Culicidae). Journal of Medical Entomology, 46(3):451–459, may 2009.

21. B. Gompertz. On the nature of the function expressive of the law of human mortality, and on the new mode of determining the value of life contingencies. Philosophical Transactions of the Royal Society of London, 115(1825):513–583, 1825.

22. D. R. Guedes, M. H. Paiva, M. M. Donato, P. P. Barbosa, L. Krokovsky, S. W. Rocha, K. L. Saraiva, M. M. Crespo, T. M. Rezende, G. L. Wallau, R. M. Barbosa, C. M. Oliveira, M. A. Melo-Santos, L. Pena, M. T. Cordeiro, R. F. O. Franca, A. L. De Oliveira, C. A. Peixoto, W. S. Leal, and C. F. Ayres. Zika virus replication in the mosquito Culex quinquefasciatus in Brazil. Emerging Microbes and Infections, 6(8), 2017.

23. F. Gunay. Farkli sabit sicakliklarda Culex quinquefasciatus (Diptera: Culicidae)’un reaksiyon normu ve kalitsalliğgi. Msc thesis, Hacettepe University, 2009.

24. F. Gunay, B. Alten, and E. D. Ozsoy. Estimating reaction norms for predictive population parameters, age specific mortality, and mean longevity in temperature-dependent cohorts of Culex quinquefasciatus Say (Diptera: Culicidae). Journal of Vector Ecology, 35(2):354–362, 2010.

25. W. S. C. Gurney, R. M. Nisbet, and J. H. Lawton. The Systematic Formulation of Tractable Single-Species Population Models Incorporating Age Structure. Journal of Animal Ecology, 52(2):479–495, 1983.

26. W. Hastings. Monte Carlo sampling methods using Markov chains and their applications. Biometrika, 57(1):97, apr 1970.

27. M. Iannelli and F. Milner. The Basic Approach to Age-structured Population Dynamics Models, Methods and Numerics. 2017.

28. A. M. Kakde, K. Patel, and S. Tayade. Role of Life Table in Insect Pest Management–A Review. IOSR Journal of Agriculture and Veterinary Science, 7(1):40–43, 2014.

29. H. Kettle and D. Nutter. StagePop: Modelling stage-structured populations in R. Methods in Ecology and Evolution, 6(12):1484–1490, 2015.

30. P. Kirk, T. Thorne, and M. Stumpf. Model selection in systems and synthetic biology. Current Opinion in Biotechnology, 2013.

31. L. P. Lefkovitch. The Study of Population Growth in Organisms Grouped by Stages. Biometrics, 21(1):1–18, 1965.

32. P. H. Leslie. On the Use of Matrices in Certain Population Mathematics. Biometrika, 33(3):183–212, 1945.

33. J. Liepe, C. Barnes, E. Cule, K. Erguler, P. Kirk, T. Toni, and M. P. H. Stumpf. ABC-SysBio-approximate bayesian computation in python with GPU support. Bioinformatics, 26(14):1797–1799, 2010.

34. J. Liepe, P. Kirk, S. Filippi, T. Toni, C. P. Barnes, and M. P. H. Stumpf. A framework for parameter estimation and model selection from experimental data in systems biology using approximate Bayesian computation. Nature Protocols, 9(2):439–456, feb 2014.

35. P. Marjoram, J. Molitor, V. Plagnol, and S. Tavaré. Markov chain Monte Carlo without likelihoods. Proceedings of the National Academy of Sciences of the United States of America, 100(26):15324–15328, 2003.

36. R. May. Uses and Abuses of Mathematics in Biology. Science, 303(5659):790, feb 2004.

37. R. Menard. Whole-genome analysis of plasmodium spp. Utilizing a new agilent technologies DNA microarray platform, volume 923. Springer, 2013.

38. S. E. Naranjo and P. C. Ellsworth. The contribution of conservation biological control to integrated control of Bemisia tabaci in cotton. Biological Control, 51(3):458–470, 2009.

39. R. M. Nisbet and W. S. Gurney. The systematic formulation of population models for insects with dynamically varying instar duration. Theoretical Population Biology, 23(1):114–135, 1983.

40. G. R. A. Okogun. Life-table analysis of Anopheles malaria vectors: Generational mortality as tool in mosquito vector abundance and control studies. Journal of Vector Borne Diseases, 42(2):45–53, 2005.

41. J. A. Patz, W. J. Martens, D. A. Focks, and T. H. Jetten. Dengue fever epidemic potential as projected by general circulation models of global climate change. Environmental Health Perspectives, 106(3):147–153, 1998.

42. G. Rosen. Time delays produced by essential nonlinearities in population growth models. Bulletin of Mathematical Biology, 49(2):253–255, 1987.

43. R. Ross. Report on the prevention of malaria in Mauritius. 1908.

44. H. Selina S. Application of Life-History Theory and Population Model Analysis to Turtle Conservation Author (s): Selina S. Heppell Published by : American Society of Ichthyologists and Herpetologists (ASIH) Stable URL : https://www.jstor.org/stable/1447430REFERE. Copeia, 1998(2):367–375, 1998.

45. D. L. Smith, K. E. Battle, S. I. Hay, C. M. Barker, T. W. Scott, and F. E. McKenzie. Ross, macdonald, and a theory for the dynamics and control of mosquito-transmitted pathogens. PLoS Pathog, 8(4):e1002588, jan 2012.

46. R. Subra. Biology and control of Culex pipiens quinquefasciatus Say, 1823 (Diptera, Culicidae) with special reference to Africa. International Journal of Tropical Insect Science, 1(04):319–338, 1981.

47. H.-J. Teng and C. Apperson. Development and survival of immature Aedes albopictus and Aedes triseriatus (Diptera: Culicidae) in the laboratory: effects of density, food, and competition on response to temperature. Journal of Medical Entomology, 37(1):40–52, 2000.

48. T. Toni and M. P. H. Stumpf. Simulation-based model selection for dynamical systems in systems and population biology. Bioinformatics, 26(1):104–110, jan 2010.

49. T. Toni, D. Welch, N. Strelkowa, A. Ipsen, and M. P. H. Stumpf. Approximate Bayesian computation scheme for parameter inference and model selection in dynamical systems. Journal of Royal Society Interface, 6(31):187–202, dec 2008.

50. A. Tran, G. L’ambert, G. Lacour, R. Beno [ineq] it, M. Demarchi, M. Cros, P. Cailly, M. Aubry-Kientz, T. Balenghien, and P. Ezanno. A rainfall- and temperature-driven abundance model for Aedes albopictus populations. IJERPH, 10(5):1698–1719, may 2013.

51. L. Xiushan, Z. Yalin, L. Youqing, and S. Josef. Life history, life table, habitat, and conservation of Byasa impediens (Lepidoptera : Papilionidae). Acta Ecologica Sinica, 26(10), 2006.

